# EV-Checklist: A fast documentation tool for enhancing transparency and accessibility in extracellular vesicle research

**DOI:** 10.1101/2024.05.17.594642

**Authors:** Rodolphe Poupardin, Martin Wolf, Nicole Maeding, Gregor Fuhrmann, Dirk Strunk

## Abstract

Extracellular vesicle (EV) research has experienced rapid growth in the past decade. While community-driven efforts, such as the MISEV guidelines (2014, 2018, 2023), have aimed to standardize reporting and enhance experimental rigor, these have not yet been universally adopted in daily EV research practices. The complexity and time commitment required by existing reporting tools can deter researchers from fully embracing them, leading to incomplete or inconsistent documentation in EV studies. Therefore, we created EV-checklist, a complementary digital tool to streamline documentation during manuscript submission and increase transparency for the reviewers, editors and readers. Our tool guides researchers through a straightforward checklist covering nomenclature, source, isolation, characterization, and functional studies of EVs. It generates a concise two-page PDF formatted table for easy integration into manuscripts and provides editors, reviewers, and readers with a clear overview of the conducted studies. EV-Checklist intends to complement existing comprehensive registries as a ‘fast-and-easy’ tool to enhance clarity and accessibility of methodological and functional insights. It may also promote higher adherence to documentation standards and in EV studies. Our tool is available on https://ev-zone.org.

## Introduction

Extracellular vesicles (EVs) have emerged as a research subject due to their roles in intercellular communication and their potential as biomarkers and therapeutic agents (Yáñez-Mó *et al*., 2015; Van Niel, D’Angelo and Raposo, 2018; Kalluri and LeBleu, 2020; Nguyen *et al*., 2020; Berumen Sánchez *et al*., 2021). A survey of publications indexed in PubMed from 2012 to 2021 (Poupardin, Wolf and Strunk, 2021) revealed that the EV research domain surged at an average annual growth rate of approximately 25.99%, leaping from 1,084 in 2012 to 8,627 by 2021, while the overall growth rate of total publications hovered around 4.38% annually. As a comparison, the iPSC field, although growing robustly, expanded at a rate of 9.67% per annum, growing from 2,131 in 2012 to 4,829 in 2021. (Figure 1). With this increasing volume of literature, fast, simple and standardized summaries are vital to ensure reproducibility and transparency of findings. The establishment of the Minimal Information for Studies of Extracellular Vesicles (MISEV) guidelines marked a milestone, advocating for rigorous reporting to enhance transparency and comparability across studies in this domain (Lötvall *et al*., 2014; Théry *et al*., 2018; Welsh *et al*., 2024).

**Figure 1:**
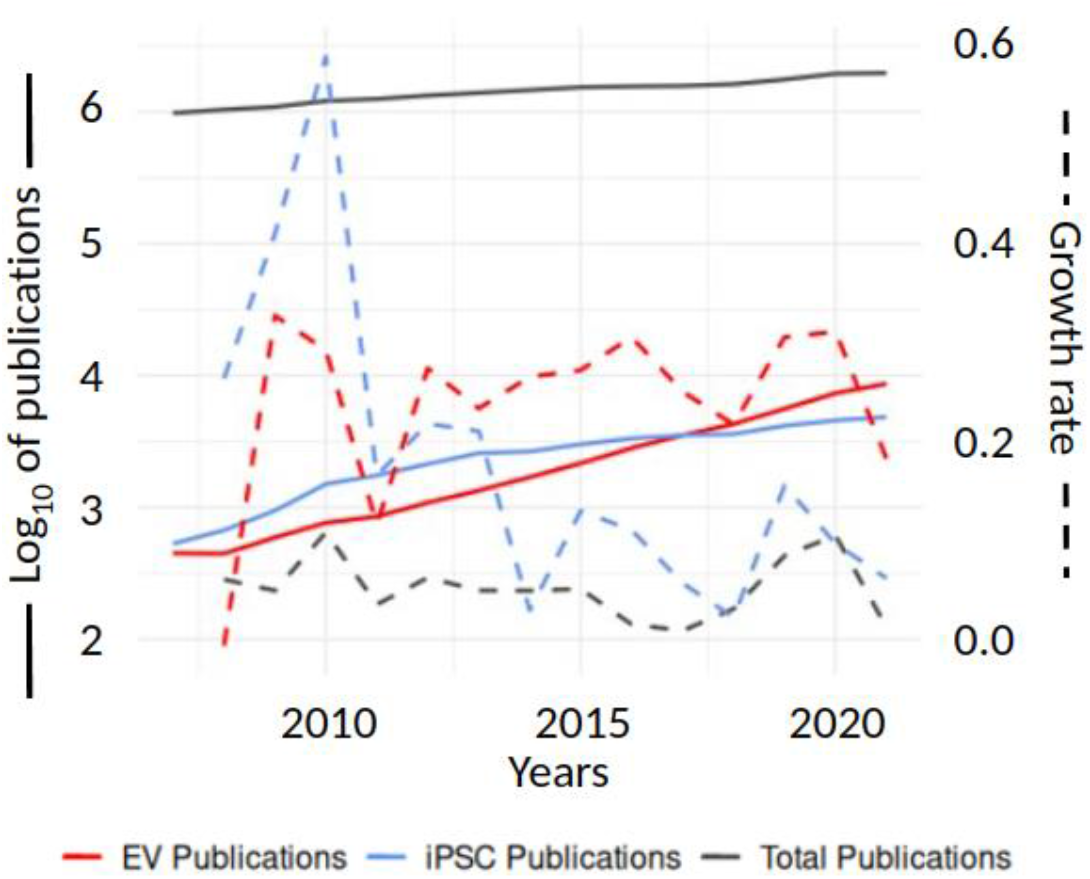
**Annual** number of PubMed publications from 2007 to 2021, illustrating the growth of publications in various scientific fields. The solid lines represent the logarithm of the total number of scientific publications (blue), publications related to extracellular vesicles and exosomes (red), and studies on induced pluripotent stem cells (iPSCs) (black). The dashed lines indicate the respective average annual growth rates for these categories, using the same color coding. The growth rate for each year is calculated as the relative increase or decrease in publication count from the previous year. The primary y-axis (left) displays the log-transformed publication counts to accommodate the wide range of values, while the secondary y-axis (right) shows the growth rates, emphasizing the dynamic nature of scientific research and literature expansion. This dual-axis plot provides a clear visual representation of both absolute publication numbers and their relative year-on-year changes.

In line with these developments, new task force initiatives have been launched to promote method-specific reporting standards. MIFlowCyt-EV offers a framework for standardized reporting of extracellular vesicle flow cytometry experiments. Similarly, MIBlood-EV (Lucien *et al*., 2023) most recently focused on enhancing transparency in reporting pre-analytical parameters for blood collection, quality control of plasma and serum parameters, processing, and storage. It was developed in response to surveys from ISEV members and the Blood EV Task Force. The tool aims to improve biobank quality, facilitate sample exchange between biobanks and labs, and enhance comparative EV studies and peer review processes. Initially provided as a user-friendly document, there is the possibility to upload the filled forms to a centralized database, though restricted to MIBlood-EV checklists only. Building upon these efforts, the International Society for Extracellular Vesicles (ISEV) has extended its guidance to cerebrospinal fluid (CSF) EV studies with the publication by (Sandau *et al*., 2024). This publication introduces comprehensive recommendations for the reproducibility of CSF EV studies and provides a detailed checklist template. This template is designed for researchers to report on key methods and some results, which can be inserted alongside the manuscript during submission, ensuring a standardized approach to documenting CSF EV research practices. However, there is no database provided for researchers to upload their CSF checklists.

In addition to these targeted initiatives, significant strides have been made using surveys (Royo *et al*., 2020) and large-scale text mining approaches to systematically analyze and understand the methodologies employed in the field (Poupardin, Wolf and Strunk, 2021; Poupardin *et al*., 2024). They serve as an invaluable resource for identifying gaps in method reporting, guiding the development of tools and guidelines that address these areas (Poupardin et al. 2024).

These initiatives encourage researchers to provide detailed methods and results reporting, contributing to the field’s overall rigor and data reproducibility. Such targeted guidelines complement broader platforms like the EV-Track https://evtrack.org/ (Van Deun *et al*., 2017) that has been instrumental in promoting comprehensive documentation, thereby contributing to elevation of transparency and reproducibility standards in EV research. Based on the amount of data reported, an EV-metric score is provided to motivate users to reach the highest standards. Additionally, data can be uploaded after publication. This database provides the currently most complete reporting system. The advantage of detailed data submission to EV-Track by reporting each EV preparation separately requires a time-intense submission process. The current lack of incentives for uploading to EV-Track could be seen as an area for improvement. The team behind EV-Track is actively working to streamline the data submission process. The anticipated release of a more user-friendly version of EV-Track reflects a commitment to continuous improvement ensuring that the platform remains at the forefront of facilitating cutting-edge research in EV studies. Other tools like ExoCarta (www.exocarta.org) and Vesiclepedia (www.microvesicles.org) also offer platforms for sharing and comparing EV data and enriching the community’s collective understanding.

A streamlined time-efficient documentation tool that covers all fields of EV research at once would be beneficial during the manuscript submission process and as a concise methods/data overview. Initiatives such as Cell Press STAR Methods or Nature’s Reporting Summary have promoted structured, transparent, and accessible reporting in life sciences. Both provide a structured format for authors to detail their experimental procedures, aiming to enhance transparency, reproducibility, and accessibility of research findings (Tonzani and Fiorani, 2021). Recognizing this, we developed a specific checklist to facilitate a swift documentation process covering important aspects of EV research, aligned with the MISEV guidelines. We applied a preliminary version of this checklist in several of our recent manuscript (Andrade *et al*., 2021; Binder *et al*., 2021; Poupardin, Wolf and Strunk, 2021; Gomes *et al*., 2022; Wolf *et al*., 2022). Based on this experience, we wanted to make our checklist publicly available and created an online tool. Upon completion, this tool generates a two-page PDF-formatted table upon completion and allows users providing users a fast overview about already completed and still pending key characterizations. Finally, this PDF could be included in manuscripts during submission offering a clear reporting of methods and quality controls.

## EV-Checklist for users: Simplicity and ease of use

The checklist serves as a concise overview of the methodologies employed and the most important characterization results, rather than delineating the specific stepwise procedures utilized for each individual sample (Figure 2). We divided the checklist into six categories: General information, source, isolation, characterization, cargo and function(s). Additionally, two optional sections were included: an “other information” field where users can provide supplementary details relevant to the study, and a feedback section, only visible to the website administrator, where users can offer suggestions for improving EV-Checklist. When initially accessing the tool, users are prompted to either generate a new unique code or enter a previously generated code. This code acts as an identifier for their checklist, enabling the save-and-resume functionality, which is crucial for accommodating the iterative nature of research and manuscript preparation.

**Figure 2:**
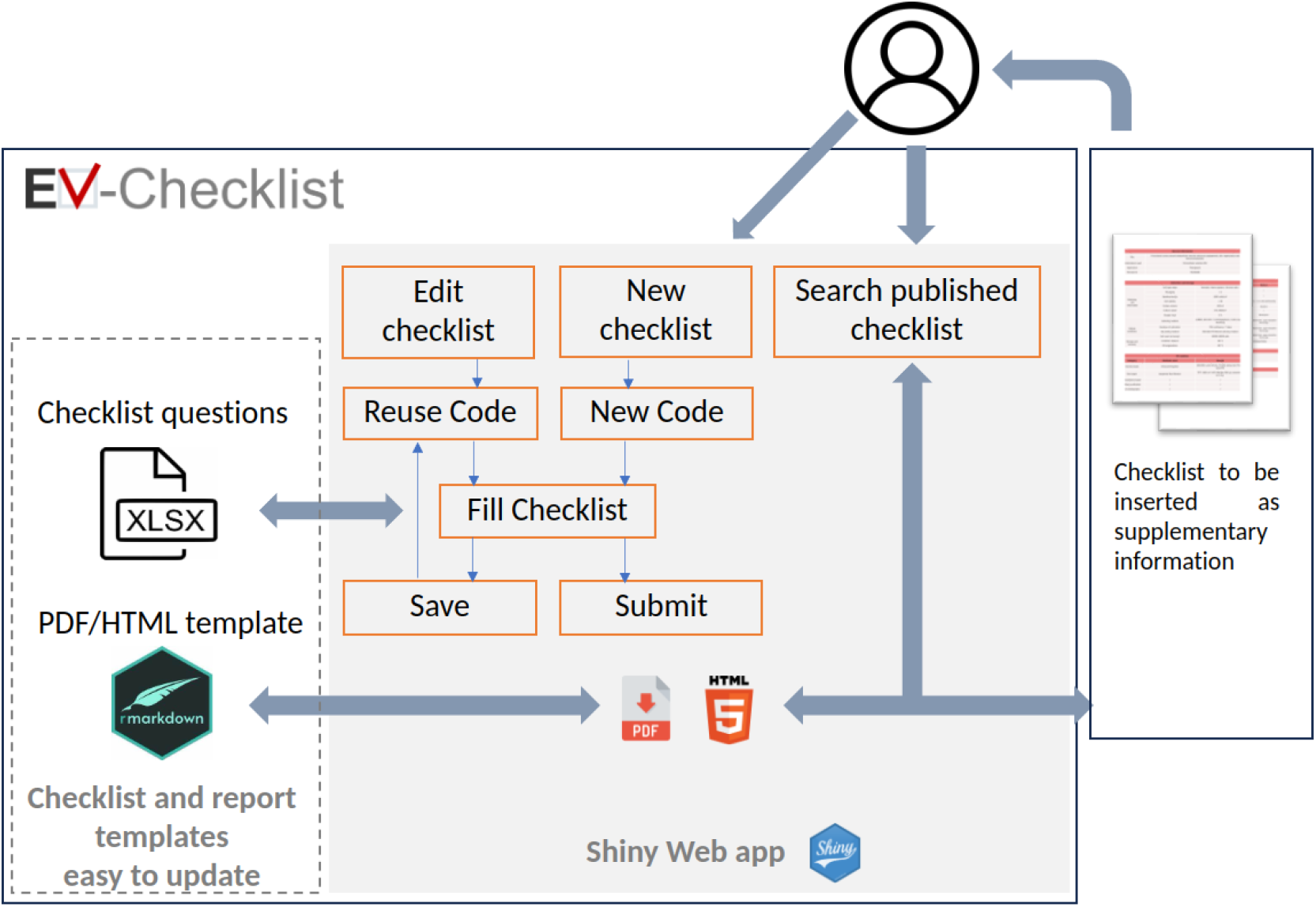
Architecture of the EV-Checklist Online Tool. Users can initiate a new checklist by obtaining a new code or continue/resubmitting one using a previously acquired code. Upon submission of a checklist, users receive a neatly formatted PDF and HTML checklist, which can be seamlessly inserted into their manuscripts. For improved visibility and transparency, all published checklists are made accessible in the checklist database. The questionnaire and PDF template are housed within an Excel file and RMarkdown document respectively, facilitating quick and effortless updates to the tool.

We organized the checklist into multiple tabs and boxes to facilitate easy navigation and to logically group related questions. Questions within the checklist are dynamically displayed based on the user’s responses to previous questions, enabling a tailored flow through the checklist. Mandatory questions are marked with an asterisk, prompting users to complete these before submission, thereby ensuring the comprehensiveness and integrity of the documentation. We optimized the application for intuitive navigation, with a clear demarcation between completed, pending, and optional fields, and aiding users in tracking their progress through the checklist. With all information at hand, it takes 15 to 25 minutes to fill out the checklist. The EV-checklist output is provided as a HTML or PDF summary table in a format ensuring a high quality, printable output that meets standards for publication (Figure 3).

**Figure 3:**
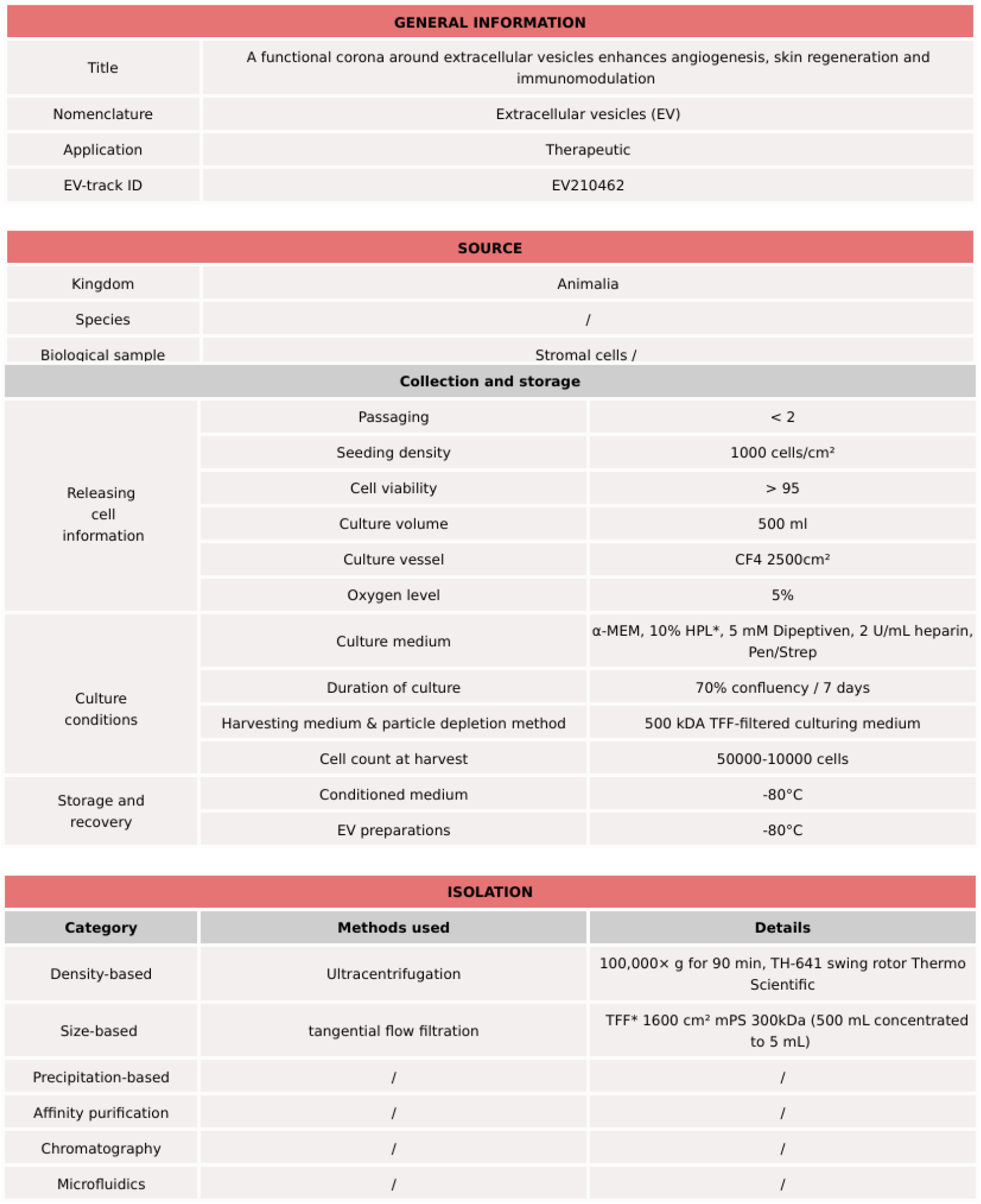
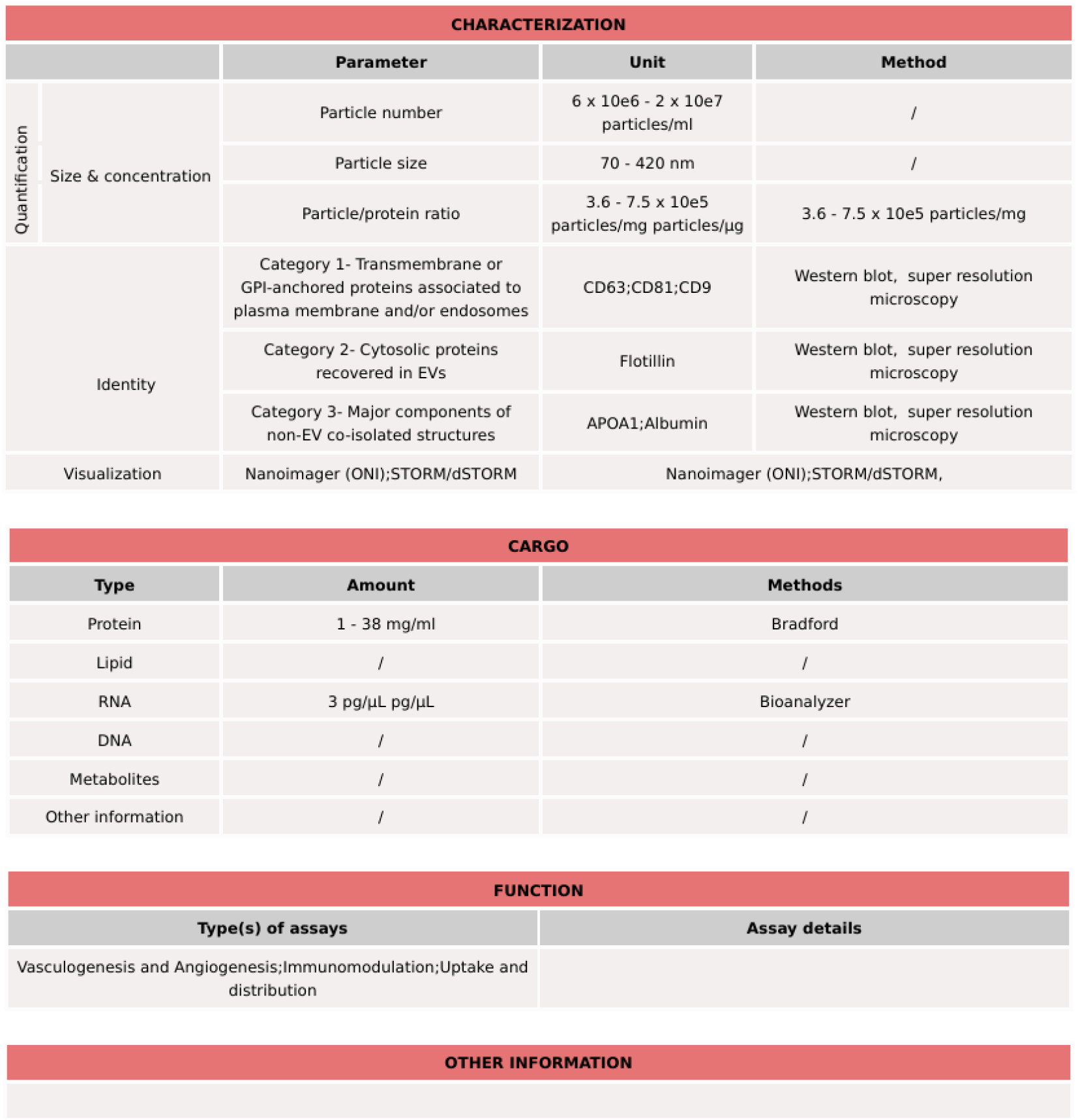
Example of EV-Checklist PDF output. The checklist is divided into six categories: General information, collection and storage, EV isolation, EV characterization, EV function(s). An optional “Other information” is also available for the submitter to provide any other relevant information.

An integral feature of our tool is the Checklist Database, which serves as a repository of all submitted checklists marked as published. This database is accessible through a dedicated tab within the tool, presenting a structured view of the diverse studies conducted in the EV field, promoting visibility of published works, and facilitating a collaborative and open research environment.

## EV-Checklist for developers: A tool optimized to be future proof

Utilizing the Shiny framework in R programming language, known for creating userfriendly interfaces while maintaining a robust backend, we developed a highly interactive web application. The checklist structure is encapsulated in a structured Excel file, which serves as a dynamic template for the checklist rendered within the tool. This design choice offers several advantages: First, the Excel file allows for easy modifications and additions to the checklist by the administrator without the necessity for code alterations. New questions can be added, or existing ones can be modified by updating the excel file. Second, the Excel file defines the types of input fields associated with each question, supporting various types of questions including text fields, radio buttons, and checkboxes. This flexibility accommodates a variety of data inputs, aiding in capturing precise and structured responses from the users. Furthermore, within the Excel file, questions are flagged as either required or optional, ensuring that critical information is not omitted and guiding users to provide all necessary details before submission. The Excel file also supports the definition of dependencies between questions. Some questions may only be relevant based on the responses to previous questions. Questions are grouped into boxes and tabs based on thematic or procedural similarities, facilitating intuitive navigation through the checklist and making it easier for users to complete and review their entries. This Excel-driven approach promotes a high degree of customization and adaptability, ensuring that the tool remains relevant and useful as the standards and requirements for EV research documentation evolve over time without requiring changes to the codebase.

Rendering of the summary table into HTML and PDF formats is facilitated through an R Markdown document designed to read the submitted responses from the database and dynamically generate a well-structured summary table. Utilizing the flextable package, formatting of the summary table is enhanced, offering a variety of table style options to achieve a publication-ready format. Moreover, the separation of the summary table rendering logic into a distinct R Markdown document, as opposed to hard coding within the Shiny application, promotes modularity and ease of maintenance. It allows for changes to the table formatting or content to be made directly within the R Markdown document without requiring modifications to the codebase of the web tool itself. Once generated in HTML format, the summary table is converted to a PDF document using the pagedown package.

## Further developments

Our tool’s modular design can host other questionnaires pertinent to the research community. In addition to its current capabilities, our tool is designed to seamlessly incorporate specialized checklists from various task force initiatives, such as MIBlood-EV (Lucien *et al*., 2023). For instance, when a user selects ‘blood’ as a sample source, the tool dynamically will integrate specific questions pertinent to blood EV research, following the MIBlood-EV guidelines. Thereby, it can easily be adapted to future advancements and to serve a broader spectrum of informational and data collection needs. Moreover, with the forthcoming release of EV-TRACK 2.0, we envisage integrating a feature that allows for the automatic import of data from EV-Track. This would enable users who have previously filled out the EV-Track questionnaire to pre-fill the checklist on our tool effortlessly, reduce redundancy and streamline the data entry process. Such integration aims to foster a more interconnected ecosystem of data sharing and retrieval.

## Conclusion

We have developed a comprehensive tool to facilitate the documentation and dissemination of experimental details in EV research. Our tool offers a user-centric interface to navigate through a structured checklist, ensuring thorough documentation of research methodologies and findings. The dynamic nature of the checklist, alongside the automated summary table generation, is a step towards reducing the documentation burden on researchers. Additionally, the feature of a publicly accessible checklist database further augments the transparency and visibility of published data. With prospects of hosting other community-relevant questionnaires and integrating with platforms like EV-Track, our tool represents a scalable and adaptable resource to foster a culture of clear and accessible reporting within the EV research landscape.

## Acknowledgments

We would like to thank our beta testers for helping us to improve EV-Checklist: André Görgens, André Cronemberger Andrade, Fausto Gueths Gomes, Paolo Bergese, Wolfgang Holnthoner, Johannes Österreicher, Marko Morávek, Nedim Haciosmanoglu, Fatih Inci, Alexander Otahal and Andrea de Luna.

## References

Andrade, A.C. et al. (2021) ‘Hypoxic conditions promote the angiogenic potential of human induced pluripotent stem cell-derived extracellular vesicles’, International Journal of Molecular Sciences, 22(8). Available at: 10.3390/ijms22083890.

Berumen Sánchez, G. et al. (2021) ‘Extracellular vesicles: mediators of intercellular communication in tissue injury and disease’, Cell Communication and Signaling 2021 19:1, 19(1), pp. 1–18. Available at: 10.1186/S12964-021-00787-Y.

Binder, H.M. et al. (2021) ‘Scalable enrichment of immunomodulatory human acute myeloid leukemia cell line-derived extracellular vesicles’, Cells, 10(12). Available at: 10.3390/cells10123321.

Van Deun, J. et al. (2017) ‘EV-TRACK: Transparent reporting and centralizing knowledge in extracellular vesicle research’, Nature Methods. Nature Publishing Group, pp. 228–232. Available at: 10.1038/nmeth.4185.

Gomes, F.G. et al. (2022) ‘Synergy of Human Platelet-Derived Extracellular Vesicles with Secretome Proteins Promotes Regenerative Functions’, Biomedicines, 10(2). Available at: 10.3390/biomedicines10020238.

Kalluri, R. and LeBleu, V.S. (2020) ‘The biology, function, and biomedical applications of exosomes’, Science (New York, N.Y.), 367(6478). Available at: 10.1126/SCIENCE.AAU6977.

Lötvall, J. et al. (2014) ‘Minimal experimental requirements for definition of extracellular vesicles and their functions: A position statement from the International Society for Extracellular Vesicles’, Journal of Extracellular Vesicles, 3(1). Available at: 10.3402/jev.v3.26913.

Lucien, F. et al. (2023) ‘MIBlood-EV: Minimal information to enhance the quality and reproducibility of blood extracellular vesicle research’, Journal of Extracellular Vesicles, 12(12), p. 12385. Available at: 10.1002/jev2.12385.

Nguyen, V.V.T. et al. (2020) ‘Functional assays to assess the therapeutic potential of extracellular vesicles’, Journal of Extracellular Vesicles. John Wiley and Sons Inc. Available at: 10.1002/jev2.12033.

Van Niel, G., D’Angelo, G. and Raposo, G. (2018) ‘Shedding light on the cell biology of extracellular vesicles’, Nature Reviews Molecular Cell Biology, 19(4), pp. 213–228. Available at: 10.1038/nrm.2017.125.

Poupardin, R. et al. (2024) ‘Advances in Extracellular Vesicle Research Over The Past Decade: Source And Isolation Method Are Connected with Cargo And Function’, Advanced Healthcare Materials [Preprint]. Available at: 10.1002/adhm.202303941.

Poupardin, R., Wolf, M. and Strunk, D. (2021) ‘Adherence to minimal experimental requirements for defining extracellular vesicles and their functions’, Advanced Drug Delivery Reviews. Elsevier B.V. Available at: 10.1016/j.addr.2021.113872.

Royo, F. et al. (2020) ‘Methods for Separation and Characterization of Extracellular Vesicles: Results of a Worldwide Survey Performed by the ISEV Rigor and Standardization Subcommittee’, Cells, 9(9). Available at: 10.3390/CELLS9091955.

Sandau, U.S. et al. (2024) ‘Recommendations for reproducibility of cerebrospinal fluid extracellular vesicle studies’, Journal of Extracellular Vesicles, 13(1), p. 12397. Available at: 10.1002/JEV2.12397.

Théry, C. et al. (2018) ‘Minimal information for studies of extracellular vesicles 2018 (MISEV2018): a position statement of the International Society for Extracellular Vesicles and update of the MISEV2014 guidelines’, Journal of Extracellular Vesicles, 7(1). Available at: 10.1080/20013078.2018.1535750.

Tonzani, S. and Fiorani, S. (2021) ‘The STAR Methods way towards reproducibility and open science’, iScience, 24(4). Available at: 10.1016/J.ISCI.2021.102137.

Welsh, J.A. et al. (2024) ‘Minimal information for studies of extracellular vesicles (MISEV2023): From basic to advanced approaches’, Journal of Extracellular Vesicles, 13(2), p. e12404. Available at: 10.1002/jev2.12404.

Wolf, M. et al. (2022) ‘A functional corona around extracellular vesicles enhances angiogenesis, skin regeneration and immunomodulation’, Journal of Extracellular Vesicles, 11(4). Available at: 10.1002/jev2.12207.

Yáñez-Mó, M. et al. (2015) ‘Biological properties of extracellular vesicles and their physiological functions’, Journal of Extracellular Vesicles, 4(2015), pp. 1–60. Available at: 10.3402/jev.v4.27066.

